# Brown Adipose Tissue is Associated with Improved Cardiometabolic Health and Regulates Blood Pressure

**DOI:** 10.1101/2020.02.08.933754

**Authors:** Tobias Becher, Srikanth Palanisamy, Daniel J. Kramer, Sarah J. Marx, Andreas G. Wibmer, Ilaria Del Gaudio, Scott D. Butler, Caroline S. Jiang, Roger Vaughan, Heiko Schöder, Annarita Di Lorenzo, Allyn Mark, Paul Cohen

## Abstract

White fat stores excess energy, while brown and beige fat dissipate energy as heat^1^. These thermogenic adipose tissues markedly improve glucose and lipid homeostasis in mouse models, though the extent to which brown adipose tissue (BAT) influences metabolic and cardiovascular disease in humans is unclear^2, 3, 4^. Here, we categorized 139,224 ^18^F-FDG PET/CT scans from 53,475 patients by presence or absence of BAT and used propensity score matching to assemble a study cohort. Individuals with BAT showed lower prevalences of cardiometabolic diseases. Additionally, BAT independently correlated with lower odds of type II diabetes, coronary artery disease and congestive heart failure. These findings were supported by improved glucose, triglyceride and high-density lipoprotein values. The effects of BAT were more pronounced in overweight and obesity, indicating that BAT can offset the deleterious effects of obesity. Strikingly, we also found lower rates of hypertension among patients with BAT. Studies in a mouse model with genetic ablation of beige fat demonstrated elevated blood pressure due to increased sensitivity to angiotensin II in peripheral resistance arteries. In addition to highlighting a role for BAT in promoting overall cardiometabolic health, this study reveals a new link between thermogenic adipose tissue and blood pressure regulation.

## Main

As early as 2003, reports described increased uptake of the labeled glucose analogue ^18^F-fluorodeoxyglucose (^18^F-FDG) on positron emission tomography (PET) in areas corresponding to supraclavicular fat on computed tomography (CT), suggesting the presence of metabolically-active BAT in adult humans^5, 6^. This tissue has received intense interest from the biomedical community since 2009 when a series of papers confirmed the presence of active BAT in adults, which correlated with lower body mass index (BMI), decreased age, colder outdoor temperature, and female sex, as well as an association with decreased fasting glucose levels^7, 8, 9, 10, 11, 12^. Since then, small prospective studies in healthy humans have demonstrated that cold-activated BAT is associated with increased whole body energy expenditure and increased disposal of glucose and free fatty acids^13, 14, 15^. Although these properties have generated great enthusiasm for BAT as a therapeutic target for obesity and its associated diseases, these studies have been too small to definitively address whether brown fat is a clinically meaningful modulator of metabolic and cardiovascular disease in humans.

To answer this central question, we reviewed 139,224 ^18^F-FDG PET/CT reports conducted in 53,475 patients between June 2009 and March 2018 at Memorial Sloan Kettering Cancer Center (MSKCC) (**Fig. 1a**). ^18^F-FDG PET/CT was conducted for cancer diagnosis, staging, monitoring of treatment response, and surveillance, and it is protocol at MSKCC to report on BAT status in each study. BAT was reported on 7,923 (5.7%) ^18^F-FDG PET/CT scans (**Fig. 1b, Extended Data Table 1**) in 5,070 (9.5%) patients (**Fig. 1c, Extended Data Table 2**), consistent with prior studies^5, 6, 11, 16^. To assess accuracy of reporting and our search strategy, all scans conducted in 2016 with reported BAT were manually reviewed. Of the 1,139 scans that reported brown fat in 2016, 1,132 (99.4%) PET scans showed increased ^18^F-FDG uptake in regions identified as fat on CT in typical locations (neck, supraclavicular, axillary, mediastinal, paraspinal and abdominal) while 4 (0.4%) scans reported a possible signature and 3 (0.3%) scans were false-positive since they reported resolution of previously reported brown fat. While the usage of ^18^F-FDG PET/CT increased over the years in this study, the prevalence of BAT reporting remained consistent (**Extended Data Fig. 1**).

**Fig. 1:**
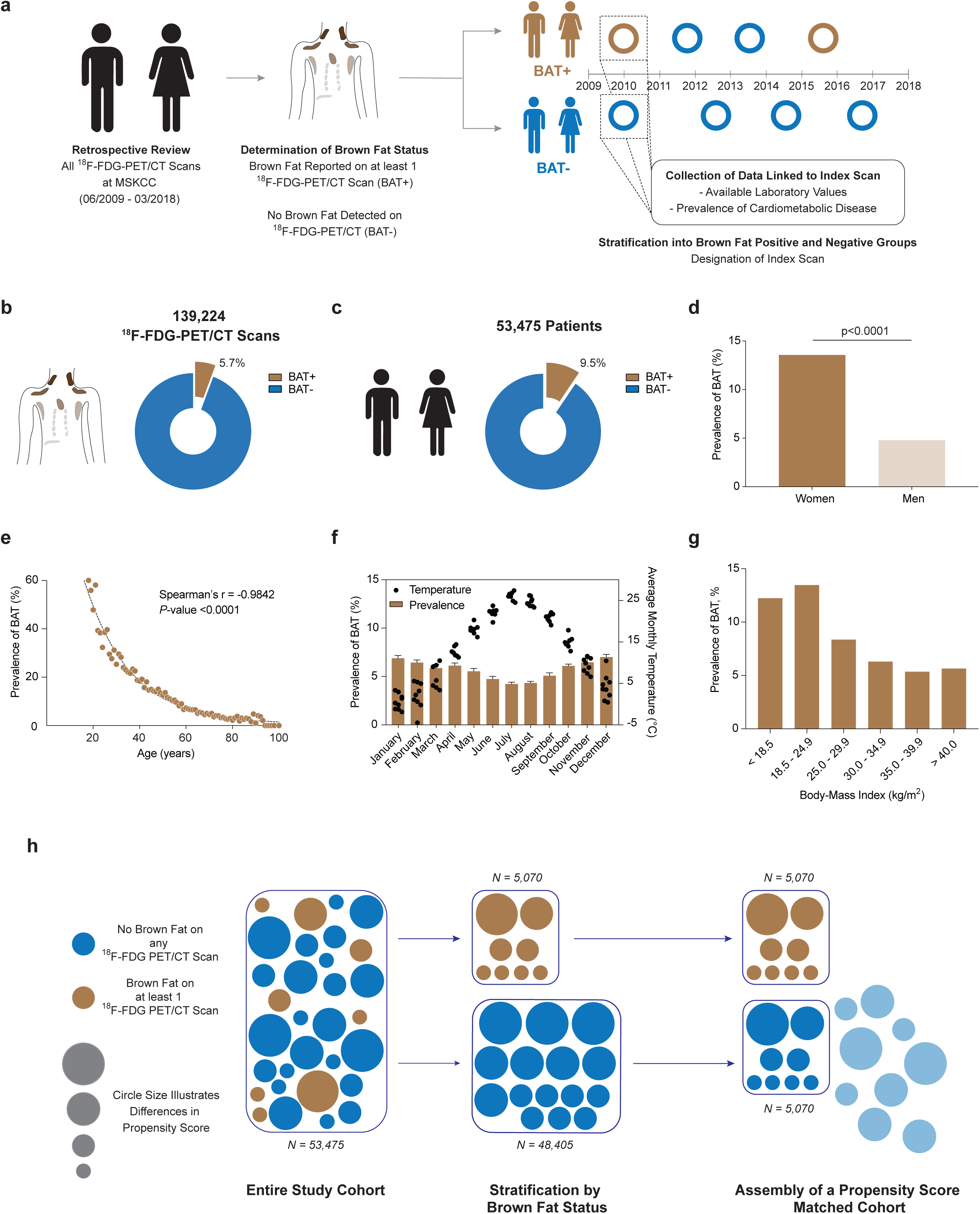
**a,** ^18^F-FDG PET/CT reports were reviewed for reporting of brown adipose tissue between June 2009 and March 2018 and individuals were stratified by presence or absence of brown fat. An index scan was assigned and diagnosis codes and available laboratory values in temporal vicinity were collected. **b**, Brown fat was reported in 5.7% of all ^18^F-FDG PET/CT scans (n=139,224). **c**, Brown fat was reported in 9.5% of all patients (n=53,475). **d**, Prevalence of brown fat was higher in females compared with males. Groups were compared by chi-square test, *P*-value is two-tailed. **e**, Brown fat prevalence was inversely correlated with age and followed a one-phase decay. Spearman’s rank correlation coefficient was calculated to assess the correlation between brown fat prevalence and age, *P*-value is two-tailed. **f**, Brown fat prevalence, based on ^18^F-FDG PET/CT scans was inversely correlated with outdoor temperature in the month of the scan between June 2009 and March 2018. Bars depict means, error bars are s.e.m., dots are mean temperature for each year between 2009 and 2018. **g,** Brown fat prevalence was inversely correlated with body mass index. **h**, Propensity score matching was used to identify a matched cohort based on age, sex, body mass index and outdoor temperature in the month of the index scan.

Patients were categorized by the presence or absence of BAT, and for those with multiple scans an index scan was assigned (**Fig. 1a**, described in methods). Brown fat was more prevalent among women (13.6 vs. 4.8%, *P*<0.0001), decreased with age (r=-0.9842, *P*<0.0001), and was inversely correlated with ambient temperature (r=-0.7093, *P*<0.0001) and BMI (r=-0.4360, *P*=0.0007), in accord with prior smaller retrospective studies (**Fig. 1d-g and Extended Data Fig. 2a and b)**^10, 11^.

Propensity score matching (PSM) allows for pairing of subjects with and without brown fat, while accounting for covariates that predict the likelihood of having brown fat. We used PSM to generate a study cohort without brown fat (*N* = 5,070), matching the individuals with brown fat based on sex, age, BMI and outdoor temperature at the time of the index scan (**Fig. 1h**). This resulting study cohort was balanced across the aforementioned variables (**Tab. 1; Extended Data Fig. 3**).

**Table 1.** Characteristics of Patients with and without Brown Fat Before and After Propensity Score Matching

| Baseline Variable | Before Propensity Score Matching |  | After Propensity Score Matching |  | P-Value * |
| --- | --- | --- | --- | --- | --- |
|  | Brown Adipose Present (BAT+) | Brown Adipose Absent (BAT-) | Brown Adipose Present (BAT+) | Brown Adipose Absent (BAT-) |  |
|  | (N=5 070) | (N=48 405) | After Matching (N=5 070) | After Matching (N=5 070) |  |
| Age, mean (SD) | 49.0 (16.3) | 61.7 (14.0) | 49.0 (16.3) | 49.2 (15.8) | 0.5863 |
| Sex, No. (%) |  |  |  |  | 1.0000 |
| Women | 3 879 (76.5) | 24 711 (51.1) | 3 879 (76.5) | 3 879 (76.5) |  |
| Men | 1 191 (23.5) | 23 694 (49.0) | 1 191 (23.5) | 1 191 (23.5) |  |
| BMI, mean (SD) | 25.8 (5.3) | 27.8 (5.9) | 25.8 (5.3) | 25.6 (5.5) | 0.0830 |
| BMI category, No. (%) |  |  |  |  | <0.0001 |
| < 25.0 | 2,564 (50.6) | 16,578 (34.3) | 2,564 (50.6) | 2,639 (52.1) |  |
| 25.0 – 29.9 | 1,589 (31.3) | 17,436 (36.0) | 1,589 (31.3) | 1,491 (29.4) |  |
| > 30.0 | 917 (18.1) | 14,391 (29.7) | 917 (18.1) | 940 (18.5) |  |
| Race, No. (%) |  |  |  |  | <0.0001 |
| Asian | 402 (7.9) | 2 851 (5.9) | 402 (7.9) | 414 (8.2) |  |
| African American | 561 (11.1) | 3 396 (7.0) | 561 (11.1) | 434 (8.6) |  |
| Caucasian | 3 694 (72.9) | 39 225 (81.0) | 3 694 (72.9) | 3 862 (76.2) |  |
| Other or Unknown | 413 (8.2) | 2 933 (6.1) | 413 (8.2) | 360 (7.1) |  |
| Ethnicity, No. (%) |  |  |  |  | 0.0281 |
| Hispanic | 398 (7.9) | 2 803 (5.8) | 398 (7.9) | 330 (6.5) |  |
| Non-Hispanic | 4 458 (87.9) | 43 059 (89.0) | 4 458 (87.9) | 4 534 (89.4) |  |
| Unknown | 214 (4.2) | 2 543 (5.3) | 214 (4.2) | 206 (4.1) |  |
| Cardiovascular Risk Factors, No. (%) |  |  |  |  |  |
| Family History of Cardiovascular Disease | 1 029 (20.3) | 11 714 (24.2) | 1 029 (20.3) | 1 086 (21.4) | 0.1636 |
| Smoking Status |  |  |  |  | <0.0001 |
| Current Smoker | 513 (10.1) | 5 390 (11.1) | 513 (10.1) | 588 (11.6) |  |
| Previous Smoker | 1 169 (23.1) | 17 755 (36.7) | 1 169 (23.1) | 1 261 (24.9) |  |
| Never Smoker | 2 999 (59.2) | 19 923 (41.2) | 2 999 (59.2) | 2 654 (52.4) |  |
| Unknown | 389 (7.7) | 5 337 (11.0) | 389 (7.7) | 567 (11.2) |  |
| 18F-FDG PET/CT Characteristics |  |  |  |  |  |
| Total Number of Scans per Patient, n (%) | 4.1 (4.0) | 2.5 (2.6) | 4.1 (4.0) | 2.5 (3.8) | <0.0001 |
| Temperature (°C Month of Reference Scan), mean (SD) | 11.9 (8.6) | 13.6 (8.8) | 11.9 (8.6) | 12.0 (8.8) | 0.7242 |
| Beta blocker, n (%) | 656 (12.9) | 12 569 (26.0) | 656 (12.9) | 899 (17.7) | <0.0001 |

\*P-value for the comparison of BAT+ and BAT- individuals in the propensity score matched group.
BAT, brown adipose tissue. SD, standard deviation. BMI, body mass index.

Individuals with brown fat showed significantly lower prevalence of type II diabetes (4.6 vs. 9.1%, *P*<0.0001) and dyslipidemia (18.9 vs. 20.6%, *P*=0.0318). This protective effect was also seen for cardiovascular disease, including atrial fibrillation/flutter (2.8 vs. 3.7%, *P*=0.0071), coronary artery disease (3.1 vs. 5.0%, *P*<0.0001), cerebrovascular disease (2.1 vs. 3.1%, *P*=0.0011), congestive heart failure (1.0 vs. 2.1%, *P*<0.0001) and hypertension (26.7 vs. 29.6%, *P*=0.0003) (**Fig. 2a**).

**Fig. 2:**
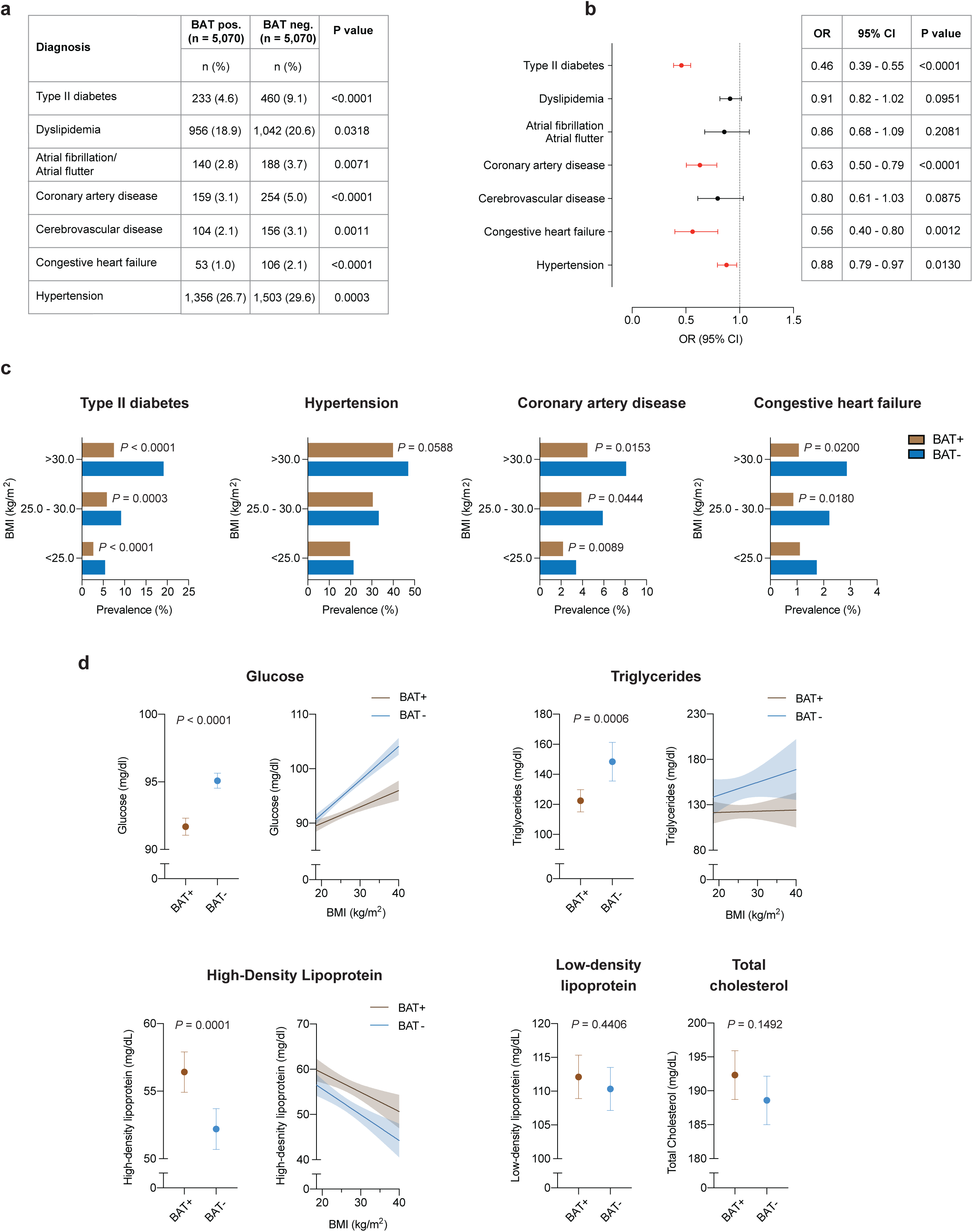
**a,** Comparison of cardiometabolic disease prevalence between individuals with and without brown adipose tissue (BAT). Groups were compared by chi-square test, all *P*-values are two-sided. **b**, Forrest plots illustrate the association between brown fat status and cardiometabolic disease in the propensity score matched cohort. Circles and bars represent odds ratios (ORs) and 95% confidence intervals (CIs), respectively, and were calculated using multivariable logistic regression analysis with adjustment for age, sex, body mass index, outdoor temperature in the month of the index scan, ethnicity, race, smoking status, family history of cardiovascular disease, beta blocker use, cancer site and cancer stage. All *P*-values are two sided. Black circles and CI bars represent non-significant associations, red circles and CI bars represent statistically significant associations at *P* < 0.05. **c**, Prevalence of cardiometabolic disease stratified by brown status and body mass index using the WHO categories for normal and underweight (body mass index (BMI) < 25.0 kg/m^2^), overweight (BMI between 25.0 and 30.0 kg/m^2^) and obesity (BMI > 30.0 kg/m^2^). Patients per category: BMI < 25.0 kg/m^2^, with brown fat *n* = 2,564, without brown fat *n =* 2,639; BMI between 25.0 and 30.0 kg/m^2^, with brown fat *n* = 1,589, without brown fat *n =* 1,491; BMI > 30.0 kg/m^2^, with brown fat *n* = 917, without brown fat *n =* 940. Significance was assessed using multivariable logistic regression adjusted for age, sex, body mass index, outdoor temperature in the month of the index scan, ethnicity, race, smoking status, family history of cardiovascular disease, beta blocker use, cancer site and cancer stage. *P* values are depicted for values <0.1. All *P* values are two sided. **d**, Comparison of available laboratory values between matched individuals with and brown fat (glucose; with brown fat n = 5,033 (99.3%), without brown fat n = 4,963 (97.9%); triglycerides; with brown fat n = 732 (14.4%), without brown fat n = 734 (14.5%); high-density lipoprotein; with brown fat n = 596 (11.8%), without brown fat n= 578 (11.4%); low density lipoprotein; with brown fat n = 543 (10.7%), without brown fat n = 527 (10.4%); total cholesterol; with brown fat n = 637 (12.6%), without brown fat n = 630 (12.4%). Groups were compared with Student’s t-test, dots represent means, error bars 95% confidence intervals, all *P*-values are two-sided. Values with statistically significant differences were stratified by body mass index and a linear regression line for body mass index values between 18.5 and 40.0 kg/m^2^ was calculated. Shaded areas depict 95% confidence intervals.

Cancer-associated characteristics such as cachexia, cancer type and stage have previously been associated with differences in brown fat prevalence on ^18^F-FDG PET/CT, as has the use of beta-blocker drugs^11, 17, 18, 19^. These associations were also observed in our study (**Extended Data Table 3 and Table 1)**. We used logistic regression analysis, adjusted for cancer site, cancer stage, and beta-blocker use, and demographic variables (including race, ethnicity, smoking status) to determine whether brown fat independently correlated with cardiometabolic disease. Multivariable logistic regression analysis identified brown fat as an independent negative predictor of type II diabetes (OR=0.46; 95% CI=0.39-0.55; *P*<0.0001), coronary artery disease (OR=0.63; 95% CI=0.50-0.79; *P*<0.0001), congestive heart failure (OR=0.56; 95% CI=0.40-0.80; *P*=0.0012) and hypertension (OR= 0.88; 95% CI=0.79-0.97; *P*=0.0130) (**Fig. 2b and Extended Data Table 4**).

Next, we stratified these data by BMI to determine whether the association between brown fat and improved cardiometabolic health was retained in overweight and obese individuals. As the prevalence of metabolic and cardiovascular disease increased with higher BMI, this effect was mitigated in individuals with brown fat (**Fig. 2c**). The benefit associated with brown fat was most striking for type II diabetes and coronary artery disease. For example, the prevalence of type II diabetes in individuals with BMI > 30 kg/m^2^ and brown fat was less than half of the prevalence in obese individuals without brown fat (7.5 vs. 19.1%, *P*<0.0001). Significantly lower prevalences of hypertension (39.9 vs. 47.0%, *P*=0.0020) and congestive heart failure (1.1 vs. 2.9%, *P*=0.0060) were also observed in individuals with obesity and brown fat. Using the multivariable logistic regression model described above, we confirmed that brown fat was independently associated with the observed decrease in cardiometabolic disease prevalence in obese individuals (**Fig 2c**).

Next, we analyzed available laboratory values in temporal vicinity to the index scan to further characterize the metabolic profile of the cohort. We noted distinct improvements in glucose (91.7 vs. 95.1 mg/dl, *P*<0.0001), triglycerides (122.4 vs. 148.4 mg/dl, *P*=0.0006), and HDL levels (56.4 vs. 52.2 mg/dl, *P*=0.0001) in individuals with brown fat, while there were no differences for LDL (112.1 vs. 110.3 mg/dl, *P*=0.4406) and total cholesterol (192.3 vs. 188.6 mg/dl, *P*=0.1492) (**Fig. 2d and Extended Data Table 5**). By stratifying laboratory values by BMI, we found that changes in fasting blood glucose and triglycerides associated with elevated BMI were offset in individuals with brown fat, and HDL levels were higher across all BMI categories in individuals with brown fat (**Fig. 2d**). There were no apparent differences in laboratory values measuring renal function (creatinine 0.83 vs. 0.84 mg/dl, *P*=0.4537 and blood urea nitrogen 14.6 vs. 14.8 mg/dl, *P*=0.1532) and thyroid function (TSH 2.95 vs. 2.49 mIU/L, *P*=0.1764) (**Extended Data Table 5**). However, both leukocyte (6.2 vs. 6.8 10^9^ cells/L, *P*<0.0001) and platelet counts (250 vs. 260 10^9^ cells/L, *P*=0.0007) were significantly decreased in subjects with brown fat, suggesting potential roles for brown adipose beyond direct regulation of lipid and glucose metabolism (**Extended Data Table 5**). In summary, our data support an important role for brown fat in mitigating metabolic disease and its cardiovascular sequelae, particularly in obese individuals.

Thermogenic fat promotes energy expenditure, and there is increasing evidence that this tissue plays roles in metabolism beyond thermogenesis itself^20, 21, 22^. While thermogenesis clearly modulates glucose and lipid metabolism, an association between thermogenic fat and blood pressure has not been described until now. Furthermore, transcriptomic profiling indicates that murine beige fat, as opposed to developmentally pre-formed brown fat, closely approximates the inducible brown fat detected by PET/CT in adult humans^23^. We therefore used adipocyte-specific PRDM16 knockout (Adipo-PRDM16 KO) mice (**Extended Data Fig. 4a, b, and c**) with an ablation of beige fat function to investigate the potential link between thermogenic fat and blood pressure control^24^. Interestingly, a previous genome-wide association study demonstrated an association between a coding SNP in exon 9 of PRDM16 (leading to the missense mutation Pro633Leu) and hypertension in humans^25^.

Adipo-PRDM16 KO animals were implanted with radiotelemetric devices to monitor blood pressure over a three-week period (**Fig. 3a**). Our studies were conducted in 11 to 14-week-old mice on a standard chow diet, and body weight was equivalent between genotypes throughout the study (**Extended Data Fig. 4d)**. Significant increases in systolic (difference between means 3.6 mmHg; 95% CI 0.9-6.4, *P*=0.0102), diastolic (difference between means 4.2 mmHg; 95% CI 1.2-7.2, *P*=0.0071), mean arterial blood pressure (difference between means 4.0 mmHg; 95% CI 1.8–6.3, *P*=0.0007) and heart rate (difference between means 18 bpm, 95% CI 3-33, *P*=0.0202) were apparent in KO animals versus controls (**Fig. 3a and Extended Data Table 6**). After adjustment for heart rate, genotype remained independently associated with increased systolic (difference between means 2.8 mmHg; 95% CI 0.2–5.5, *P*=0.0378), diastolic (difference between means 3.3 mmHg; 95% CI 0.4–6.2, *P*=0.0281) and mean arterial blood pressure (difference between means 3.3 mmHg; 95% CI 1.0–5.6, P=0.0052) (**Extended Data Table. 7**).

**Fig. 3:**
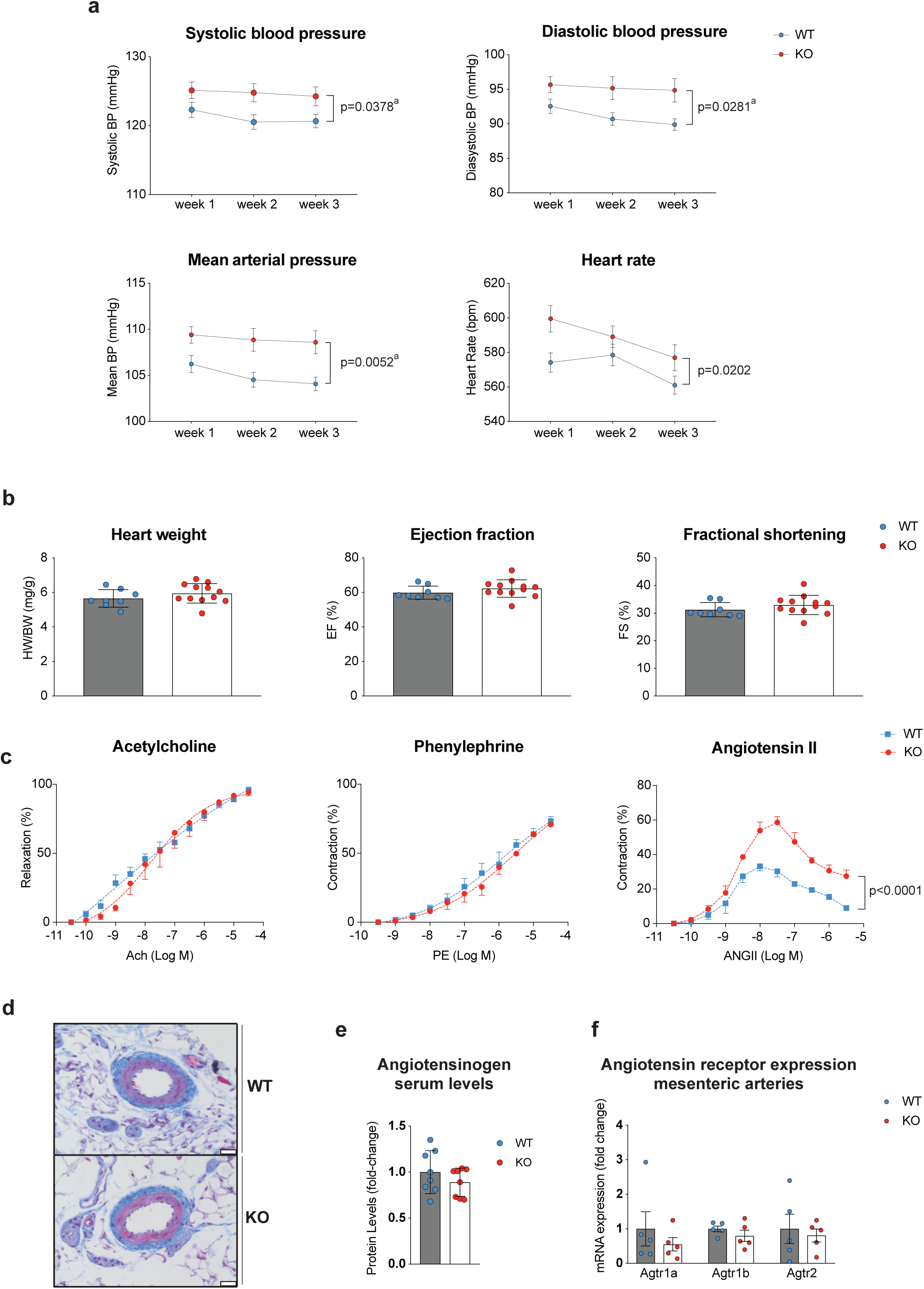
**a,** Implanted radiotelemetery devices were used to measure systolic, diastolic and mean arterial blood pressure and heart rate in freely moving mice with an adipocyte-specific knockout (KO) of *Prdm16* or wildtype (WT) controls (n=35 per group from a total of 4 independent experiments). Dots depict mean, error bars are s.e.m. A linear mixed model for repeated measures over time was used to analyze the radiotelemetry data with fixed effects of genotype (KO vs. WT), week treated as a categorical factor, and the interaction between genotype and week. ^a^ denotes *P*-values that were calculated by adjusting the above mentioned linear mixed model by heart rate to account for the observed differences in heart rate. All *P*-values are two-sided. **b**, Heart weight at necropsy normalized to body weight, ejection fraction and fractional shortening (both assessed by echocardiography) were measured in KO (n=12) and WT (n=8) animals. Bars represent means, error bars are s.d. Groups were compared by Student’s t-test. All *P*-values are two-sided. **c** Pressure myography after removal of perivascular fat was conducted in KO and WT animals (n=3 per group; 2 mesenteric arteries per mouse) to assess response to acetylcholine (endothelium-dependent vasorelaxation), phenylephrine and angiotensin II (both vasoconstrictions). Dots represent means, error bars are s.e.m. Groups were compared by two-way analysis of variance (ANOVA). All *P*-values are two-sided. **d**, Mesenteric arteries were stained with Masson’s trichrome stain to compare WT and KO animals. Representative image, n=5 per group. Scale bar is 50 µm. **e**, Angiotensinogen levels were measured in KO and WT animals using mass spectrometry (n = 8 per group). Groups were compared using Student’s t-test. All *P*-values are two-sided. **f**, Expression of angiotensin receptors on mesenteric arteries was assessed using quantitative PCR (qPCR); n=5 per group. Groups were compared using Student’s t-test. All *P*-values are two-sided.

There were no significant differences in heart weight, wall thickness, cardiac function or cardiac tissue histology, indicating that neither cardiac morphology nor function were responsible for the observed differences in systemic blood pressure (**Fig. 3b and Extended Data Fig. 5a**). We thus explored resistance vasculature morphology and function. By using the pressure myograph system, second order mesenteric arteries isolated from Adipo-PRDM16 KO mice (after dissecting off all perivascular fat) demonstrated an increased contractile response specifically to angiotensin II, with no significant differences in response to acetylcholine or phenylephrine (**Fig. 3c**). Additionally, vasodilation in response to a stepwise increase in flow was significantly decreased in Adipo-PRDM16 KO mice while myogenic tone induced by a stepwise increase in pressure did not show a difference between the genotypes (**Extended Data Fig. 5b)**. Histological analysis evidenced no apparent structural differences (**Fig. 3d**). There were no significant differences in plasma levels of angiotensinogen (**Fig. 3e**) or expression of angiotensin receptors in the mesenteric arteries (**Fig. 3f**). In summary, ablation of beige fat function in mice leads to an increased sensitivity of resistance arteries to angiotensin II, contributing to elevated systemic blood pressure.

Obesity is a major contributor to metabolic and cardiovascular disease, and 48.9% of American adults are predicted to be obese by 2030^26^. With limited effective therapies available, thermogenic fat has been implicated as a potential therapeutic target for obesity and metabolic disease^27^. In healthy adults, cold stimulation of brown fat increases energy expenditure and improves systemic glucose utilization, fatty acid oxidation, and insulin sensitivity^28, 29, 30, 31^. This observation is consistent with animal models, where transplantation of thermogenic adipose tissue mass improved glucose and triglyceride metabolism, while genetic ablation of thermogenic fat had the opposite effect^3, 32, 33^. The effects of thermogenic adipose tissue on cardiometabolic health and obesity-associated diseases in humans, however, have been largely unexplored until now.

Our study illustrates that individuals with thermogenic fat on ^18^F-FDG PET/CT have significantly improved metabolic profiles. This effect is not only limited to diabetes, but extends to coronary artery disease, congestive heart failure and hypertension. Furthermore, the effects of brown fat on metabolic and cardiovascular disease are most pronounced among individuals with elevated BMI. While obesity is generally associated with decreased brown fat function^34^, those obese individuals who retain brown fat activity appear to be protected against conditions linked to excess weight. This notion further supports the potential of brown fat as a therapeutic target beyond weight loss itself, but as a means to uncouple obesity from disease.

The reduction in the prevalence of hypertension associated with thermogenic fat was of particular interest to us, because this cannot be directly explained by improvements in glucose and lipid metabolism. Our studies in a mouse model lacking beige fat identified changes in sensitivity to angiotensin II in the peripheral resistance vasculature as a potential underlying physiological mechanism. Thermogenic fat secretes a number of endocrine factors that can affect a wide variety of tissues, and ongoing research is focused on elucidating the downstream effectors of these signals. Our work points towards a novel molecular signal linking adipose tissue phenotype to blood pressure control, and may be the first step in elucidating a mechanistic link between brown fat and hypertension.

Our study raises a number of questions regarding brown fat development and function in humans. Variability in brown fat prevalence and activity suggests a possible genetic determinant of its biology. Consistent with this, our understanding of genetics as a contributor to type II diabetes development has evolved significantly^35^. It is likely then that genetic factors contributing to brown fat development and activity may affect the pathogenesis of type II diabetes and metabolic syndrome. In our study, brown fat was associated with improved health across several organ systems. As an example, individuals without brown fat exhibited higher leukocyte counts, and elevated white blood cell counts are an established risk factor for coronary artery disease^36, 37, 38^. Decreased leukocyte counts in individuals with brown fat could, in part, explain the lower prevalence of coronary artery disease among these subjects. Our work highlights the need to determine the extent of brown fat’s effects on other organs and roles in systemic metabolism.

The major strength of our study is the size of the dataset, the largest we are aware of, and the linkage to electronic health records. This allowed us to comprehensively examine associations between brown fat and a wide variety of data. There are, however, a number of limitations that need to be acknowledged, mainly pertaining to the retrospective design and analysis of data from a cancer population. First, identification of brown fat relies on ^18^F-FDG PET/CT and consistent reporting. Our manual review of brown fat reporting over an entire year, as well as a consistent brown fat prevalence over the study period, support the reliability of reporting in this study. Furthermore, ^18^F-FDG PET/CT without prior cold stimulation tends to underestimate brown fat prevalence, which may have impacted our classification. In the absence of broadly applicable screening methods or biomarkers for brown fat, the application of ^18^F-FDG PET/CT to screen for brown fat is however still considered the gold standard. Although a number of studies have indicated associations between cancer characteristics and brown fat prevalence as measured by ^18^F-FDG PET/CT, the effect of cancer is still poorly understood. We have thus adjusted our analysis to accommodate the potential influence of cancer type and stage.

In our study, we investigated and confirmed new clinical findings linking thermogenic adipose tissue and improvements in metabolic health using an established mouse model. These findings will have the potential to enable translation of animal findings into viable human therapies. Our study highlights the therapeutic potential of modulating brown fat in humans to combat the escalating obesity crisis.

## Online Methods

### Study Design and Participants

This retrospective case-control study followed institutional guidelines and was approved by the Institutional Review Boards of The Rockefeller University and Memorial Sloan Kettering Cancer Center (MSKCC). Due to the retrospective nature of this study, the requirement for informed consent was waived. Results from this study are reported in accordance with the Strengthening the Reporting of Observational Studies in Epidemiology (STROBE) guidelines for case-control studies^39^.

We identified all patients age 18 and above who underwent ^18^F-FDG PET/CT at MSKCC from June 2009 through March 2018. This resulted in 139,224 ^18^F-FDG PET/CT scans conducted in 53,475 patients. Brown adipose is identified by increased ^18^F-FDG uptake on PET, measured in regions corresponding to adipose tissue on CT, and it is standard practice at MSKCC to document these signatures in all reports. We searched all ^18^F-FDG PET/CT reports for the terms “brown fat” or “brown adipose”. Patients were then categorized by the presence or absence of brown fat reporting. For each subject in both groups, an index scan was designated as follows: in patients with brown adipose tissue reported on at least one ^18^F-FDG PET/CT scan, the first scan that reported brown fat was defined as the index scan. In patients without brown fat on any scan, the first ^18^F-FDG PET/CT scan served as the index scan (**Fig. 1**).

### Data collection

We obtained average outdoor temperatures in New York City, measured in Central Park, for the months of scans from the U.S. National Weather Service. All patient data including demographics and self-reported race and ethnicity were collected from institutional electronic health records (EHR). Diagnoses listed up to one year after the time (month and year) of the index scan were identified using diagnostic codes from the International Classification of Diseases, 9th and 10th Revisions (ICD-9 and ICD-10) (**Supplementary Data Table 1**).

Laboratory values were recorded from the EHR if these values were first measured within three months (complete blood counts) or one year (lipid levels) of the index scan date. In case of multiple data points available, the one closest to the index scan was used for analysis. Blood glucose was routinely measured on the day of scanning. Complete blood counts from patients with hematological malignancies were excluded from analysis. With the exception of fasting blood glucose, measurement of blood values was not coupled to ^18^F-FDG PET/CT scans. Data completeness was assessed for all blood values, and available data was analyzed using complete case analysis (**Extended Data Table 5**). Data completeness ranged from 11.7% (low-density lipoproteins) to 98.1% (glucose).

### Propensity Score Matching and Identification of Study Cohort

To identify the study cohort, patients with reported brown fat were matched to patients without reported brown fat using propensity score matching. Propensity scores were estimated using a non-parsimonious multivariable logistic-regression model with brown adipose tissue status (presence or absence) as the dependent variable and age, sex, BMI, and temperature at time of scan as covariates. Matching was performed using a 1:1 protocol without replacement (greedy-matching algorithm), a caliper width of 0.2 and the variable sex was set to be exactly matched. We used standardized difference means to assess balance before and after matching (**Extended Data Fig. 3**). Matching was conducted in SAS version 9.4 (SAS Institute, Cary, NC) using the **psmatch** function.

### Outcomes

ICD-9/10 diagnoses extracted from EHRs were used to examine associations between brown fat and cardiometabolic health defined as the prevalence of type II diabetes, dyslipidemia, hypertension, coronary artery disease, congestive heart failure and cerebrovascular disease. All ICD-9/10 codes used to assign disease categories are listed in **Supplementary Data Table 1**.

### Data Analysis

We first assessed the correlation between brown fat and age, outdoor temperature in the month of the ^18^F-FDG PET/CT index scan and BMI. For age and BMI, a one-phase exponential decay curve was modeled and the correlation was assessed by calculating Spearman’s rank correlation coefficient. For temperature, a linear regression line was calculated and the correlation was assessed by calculating Spearman’s rank correlation coefficient. Prevalence of brown fat in men and women was compared using chi-squared test. Brown adipose status (either present or absent) and prevalence of cardiometabolic diseases were compared using chi-squared tests.

Next, we assessed whether brown fat status was an independent predictor of cardiovascular disease by using multivariable logistic regression analysis. The dependent variable in this analysis was the respective cardiometabolic disease and brown fat status (presence or absence) was the independent variable. To adjust for potential confounders that were selected based on previous publications, we controlled for the following variables: age, sex, BMI, outdoor temperature in Central Park in the month of the index scan, smoking status, race, ethnicity, family history of cardiovascular disease, beta blocker use, cancer site, and cancer stage.

Smoking was coded as a categorical variable with 4 levels (current smoker, previous smoker, never smoker, unknown smoking status). Race was a categorical variable with 4 levels: Caucasian, African-American, Asian and other (e.g. Pacific-Islanders, unknown). Ethnicity was a categorical variable with three levels: Hispanic, Non-Hispanic and Unknown. Cancer stage was a 6 level categorical variable consisting of Stages I-IV, not applicable (e.g. hematologic malignancies, benign tumors), and unknown cancer stage. Cancer site was recorded according to ICD 0 Site Codes and included 14 categories: C00-C14, C15-C26, C30-C39, C40-C41, C42, C43-C44, C45-C49, C50-C58, C60-C63, C64-C68, C69-C72, C73-C75, C76-C80 and other/unknown. Results were reported as odds ratio and 95% confidence interval.

Cardiometabolic diseases that were independently associated with brown fat status were further stratified by BMI categories (normal weight, defined as BMI <25km/m^2^; overweight, defined as BMI 25.0 – 30.0 kg/m^2^; obese, defined as BMI >30.0 kg/m^2^). Comparison between individuals with and without brown fat in the respective category were performed using the multivariable logistic expression model outlined above to assess whether observed differences were statistically significant. For the comparison of laboratory values, a Student’s *t*-test was performed. To compare associations between brown fat status, BMI and available lab values, a least squares linear regression line was constructed, stratified by brown status. All p-values are two-tailed, and values less than 0.05 were considered statistically significant.

### Animal Experiments

Animal care and experimentation were performed according to procedures approved by the Institutional Animal Care and Use Committee at the Rockefeller University. The Adipo-PRDM16 KO mice were generated by breeding Prdm16lox/lox mice with Adiponectin-cre mice (provided by Dr. Evan Rosen, backcrossed to a C57BL/6J background)^24^. Mice were maintained in 12 hr light:dark cycles at 23C and fed a standard irradiated rodent chow diet.

### RNA Preparation and Quantitative PCR

Total RNA was extracted from adipose tissue using TRIzol (Cat # 15596018, Thermo Fisher Scientific, Waltham, MA, USA) along with RNeasy mini kits (Cat # 74104, QIAGEN, Venlo, Netherlands). For qPCR analysis, RNA was reverse transcribed using the ABI high capacity cDNA synthesis kit (Cat # 4368813, Thermo Fisher Scientific, Waltham, MA, USA). cDNA was used in qPCR reactions containing SYBR-green fluorescent dye (Cat # 4309155, Thermo Fisher Scientific, Waltham, MA, USA). Relative mRNA expression was determined by normalization with Eukaryotic 18S ribosomal RNA levels using the DDCt method. Primer sequences are listed in **Supplementary Data Table 2.**

### Implantation of Radiotelemetry Devices and Blood Pressure Measurement

Blood pressure was measured by implantable radio transmitters as previously described^40^. Briefly, male mice at age 11-14 weeks were housed in pairs. Prior to implantation, the animals received buprenorphine (0.8 mg/Kg, s.c.) and a transmitter was implanted (model HD-X11, Data Sciences International, St. Paul, MN, USA), while mice were under isoflurane anaesthesia. The catheter tip was positioned in the thoracic aorta via the left common carotid artery and the body of the probe was inserted subcutaneously in the dorsal right flank. Animals were allowed to recover for 5–7 days. Heart rate (HR), systolic blood pressure (SBP), diastolic blood pressure (DBP), and mean arterial pressure (MAP) were collected weekly over a 24h period in freely moving conscious mice in their cages. Pressure traces were checked and animals with damping of the haemodynamic profile were excluded from the analysis

### Analysis of Blood Pressure Readings

For each mouse, systolic, diastolic and mean arterial blood pressure as well as heart rate was recorded. The mean for each parameter was calculated over a 24h period. Data from 4 experiments were pooled and used for the final analysis. A linear mixed model for repeated measures over time (SAS Proc Mixed) was used to analyze the radiotelemetry data with fixed effects of genotype (KO vs. WT), week treated as a categorical factor, and the interaction between genotype and week. This method prevented list-wise deletion due to missing data. Unstructured covariance structure was chosen with the lowest corrected Akaike’s information criteria and Bayesian information criteria. Heart rate was included as a covariate for the blood pressure models and the adjusted mean difference between KO and WT mice with 95 % confidence intervals were reported.

### Mass Spectrometry

12 µl of serum sample were incubated with 12 µL of 20mM dithiothreitol (DTT) and 16 M urea (final 8M urea, 10 mM DTT, 50mM ammonium bicarbonate (ABC)) for 1h at room temperature (RT). Another 6 µL of 50mM DTT were added to the samples (in 50 mM ABC, no urea) to ensure complete reduction. The samples were then alkylated by addition of 20 µL of 75mM iodoacetamide (IAA; 30mM final in ABC) in the dark at RT for 1.5h. 6 µg of LysC was added to each of the sample. The samples were then allowed to digest at 28° C overnight with shaking. The following day, the samples were further diluted with 60 µL of 50mM ABC. 10 µg of trypsin was added to each of the samples and samples were digested with trypsin for 6 hours at 28° C at 1400 rpm. The samples were then quenched with 10 µL of 10% TFA (pH approximately 1). 5 µL of sample was loaded onto 4 C18 empore discs. The cleaned up samples were speed vacuumed and re-dissolved in 20 µL of 0.1% LC-grade formic acid. 1 uL was loaded onto a reverse-phase nano-LC-MS/MS (EasyLC 1200, Fusion Lumos, Thermo Fisher Scientific) for analysis. Peptides were separated with a gradient in which the proportion of buffer B (H2O with 0.1% formic acid) in buffer A (acetonitrile with 0.1% formic acid) increased from 2% to 90% over 79 minutes (300 nl/min flow rate). MS and MS/MS data were recorded at resolutions of 60.000 and 30.000 with AGC’s of 1,000,000 and 50,000 respectively. Parallel reaction monitoring (PRM) was used to target peptides unique to AGT (AIQGLLVTQGGSSSQTPLLQSIVVGLFTAPGFR and LPTLLGAEANLNNIGDTNPR). PRM experiment data were analyzed with SkyLine v.4.2. Samples were analyzed in the following order: KO, WT and values were normalized to WT to derive fold-change between the two genotypes.

### Tissue Staining

Mesenteric arteries and hearts were collected from 18-week-old male mice. Briefly, mice were euthanized by isoflurane overdose and the mesenteric artery was isolated and perfused with 50 mmol/L PBS (pH 7.4) followed by 4% PFA. After fixation overnight, tissues were washed 3x with PBS and stored in ethanol 70%. Further tissue processing was performed at the Memorial Sloan Kettering Cancer Center (MSKCC) Laboratory of Comparative Pathology. Briefly, the tissues were embedded in paraffin and the dewaxed slides were first immersed in Weigert’s hematoxylin (Sigma, St. Louis, MO, U.S.A.), followed by serial staining with acid fuschin, phosphomolybdic acid and methyl blue. The tissues were then fixed in 1% acetic acid. After dehydration with increasing concentrations of alcohol, the slides were fixed in toluene, mounted in Permount and air-dried overnight before acquiring photographs.

### Echocardiography

Mice were anaesthetized with isoflurane under continuous monitoring and placed on a heating pad of a recording stage connected to a Vevo 2100 ultrasound machine. The anterior chest wall was shaved, ultrasound gel was applied and electrodes were connected to each limb to simultaneously record an electrocardiogram. Two-dimensional (short axis-guided) M-mode and B-mode measurements (at the level of the papillary muscles) were taken using an 18–32 MHz MS400. For analysis, at least three measurements were averaged; measurements within the same HR interval (450 ± 50 bpm) were used for analysis.

#### Pressure myography

Mesenteric arteries were collected from 18-week-old male mice, as described above. Second order mesenteric arteries (MA) were cleaned of surrounding fat and vascular reactivity experiments were performed as described previously ^41^. Briefly, MA were mounted on glass cannulas in a pressure myograph chamber (Danish MyoTechnology, Aarhus, Denmark) The vessel orientation in relation to the flow *in vivo* was maintained. Vessel viability was maintained using Krebs solution solution (in mM: NaCl 118, KCl 4.7, MgCl_2_ 1.2, KH_2_PO_4_ 1.2, CaCl_2_ 2.5, NaHCO_3_ 25 and glucose 10.1), at 37 °C and oxygenated (95% O_2_ and 5% CO_2_). A pressure interface controlled intraluminal pressure and flow in MA. The vessel diameter was monitored in real time using a microscope connected to a digital video camera (IC Capture) and computer software VediView 1.2 (Danish MyoTechnology, Aarhus, Denmark) with edge detection capability. MA were equilibrated for 15 min at 80 mm Hg, pre-constricted with PE (1 µM) and a cumulative concentration-response curve of Ach (0.1 nM – 30 µM) was performed to evaluate the integrity and the function of the endothelium. MA with less than 80% vasodilatory response to Ach were discarded. Vascular smooth muscle functions were evaluated by performing cumulative concentration-response curves of PE (1 nM – 30 µM), AngII (0.1 nM – 1 µM) and myogenic tone. Flow mediated vasodilation was also evaluated as previously described ^41^.

### Statistical Analysis

Unless otherwise stated, data are presented as mean ± s.e.m. and are derived from multiple experiments. Normality was assessed using the Shapiro-Wilk-test. No statistical method was used to calculate sample size, but sample size was determined based on preliminary experiments and previous publications. Analysis was carried out using unpaired Student’s *t*-test, Mann-Whitney U test or two-way ANOVA based on data distribution and as indicated. All p-values are two-tailed, and values less than 0.05 were considered statistically significant. GraphPad Prism 8 was used for statistical analysis.

## Acknowledgements

We thank Dr. Robin Davisson for guidance on establishing radiotelemetric blood pressure monitoring, Michael N. Singer and Chester Poon for expert IT support and data extraction, Rebecca Teng for assistance in accessing data, and Dr. Barry S. Coller, Dr. Andrew J. Dannenberg, Dr. Javid J. Moslehi and Melissa D. Curtis for valuable discussions during the preparation of this manuscript.

## Author Contributions

P.C. and T.B. designed and conceived the study. T.B., D.J.K, S.J.M., A.G.W., I.D.G., S.D.B. and H.S. acquired data and performed experiments. P.C., T.B., S.P., R.V., C.S.J., A.M. and A.D.L. analyzed and interpreted the data. T.B, D.J.K. and P.C. wrote the manuscript with input from all authors.

## Data Availability

Source data for figures 1, 2, 3 as well as extended data figure 1, 2, 4 and 5 are provided with the paper. All other data that support the findings of this study are available from the corresponding author upon reasonable request.

---

**Extended Data Fig. 1:**
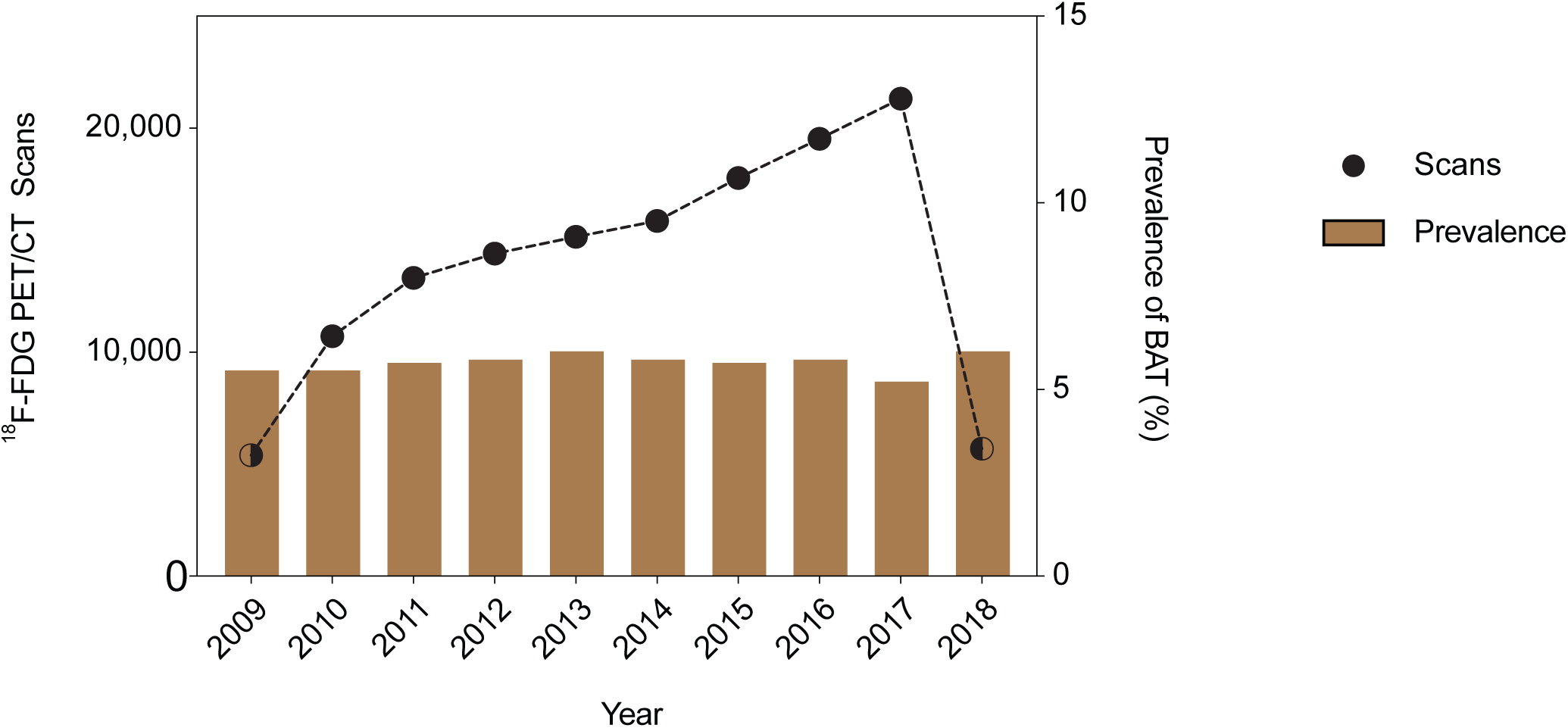
**a**, The total number of ^18^F-FDG PET/CT scans performed at MSKCC increased from 2009 to 2018 while the prevalence of reported brown fat remained stable. Dots depict total number of ^18^F-FDG PET/CT scan in each year, half filled dots illustrate years with incomplete reporting. Bars represent prevalence of ^18^F-FDG PET/CT scans with reported brown fat.

**Extended Data Fig. 2.**
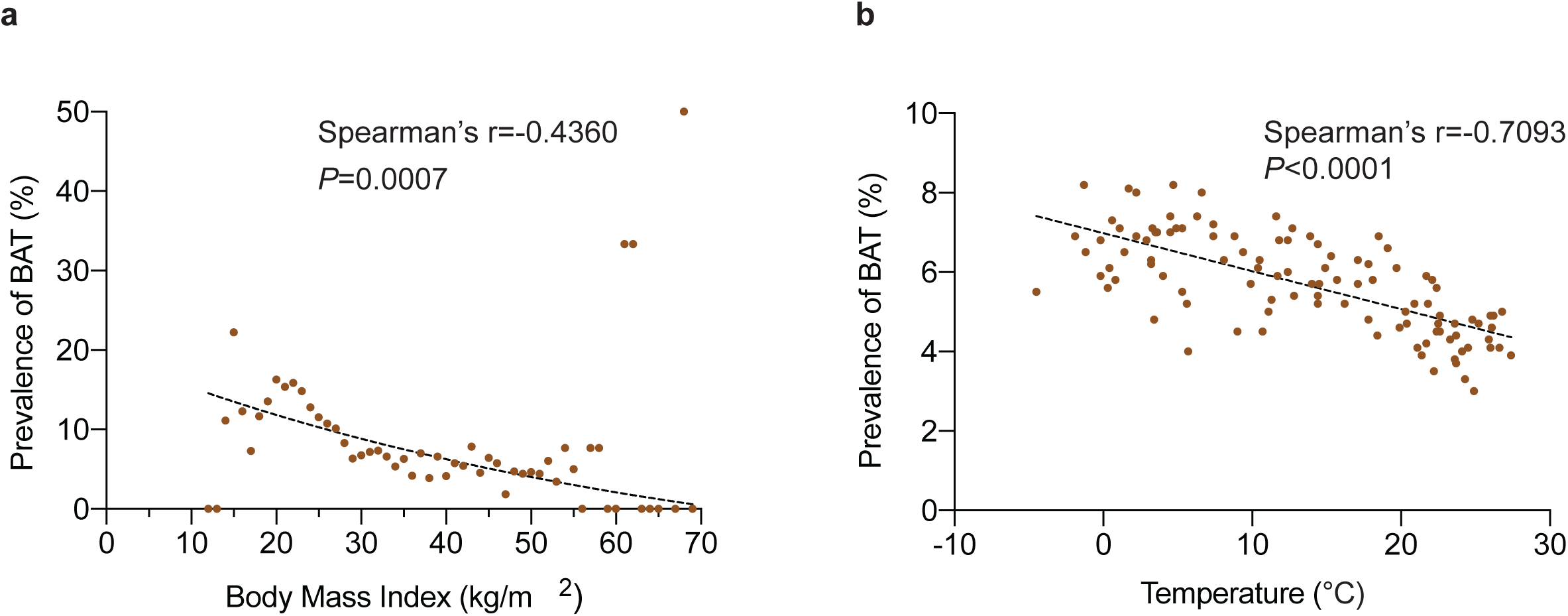
**a,** Correlation between body mass index and prevalence of brown fat reported on ^18^F-FDG PET/CT. The dotted line depicts a one-phase decay curve, *P*-value was calculated for the correlation between both variables using Spearman’s r. **b**, Correlation between outside temperature in the month of the scan and prevalence of brown fat reported on ^18^F-FDG PET/CT. The dotted line depicts a linear regression line, *P*-value was calculated for the correlation between both variables using Spearman’s r. All *P*-values are two-sided.

**Extended Data Fig. 3.**
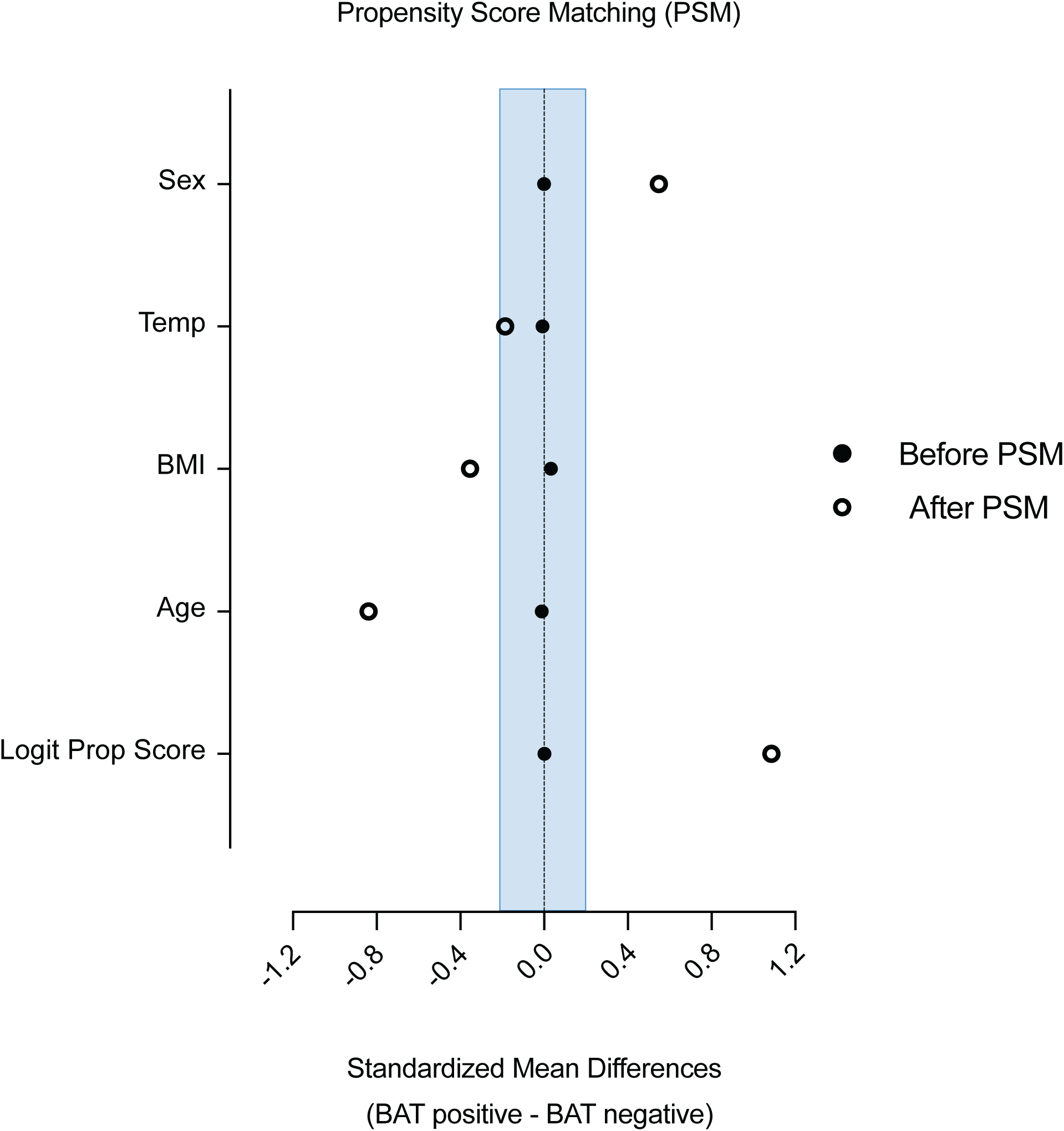
Propensity score matching was assessed by comparing standardized mean differences before and after the matching process. The blue shaded area indicates the chosen caliper width.

**Extended Data Fig. 4.**
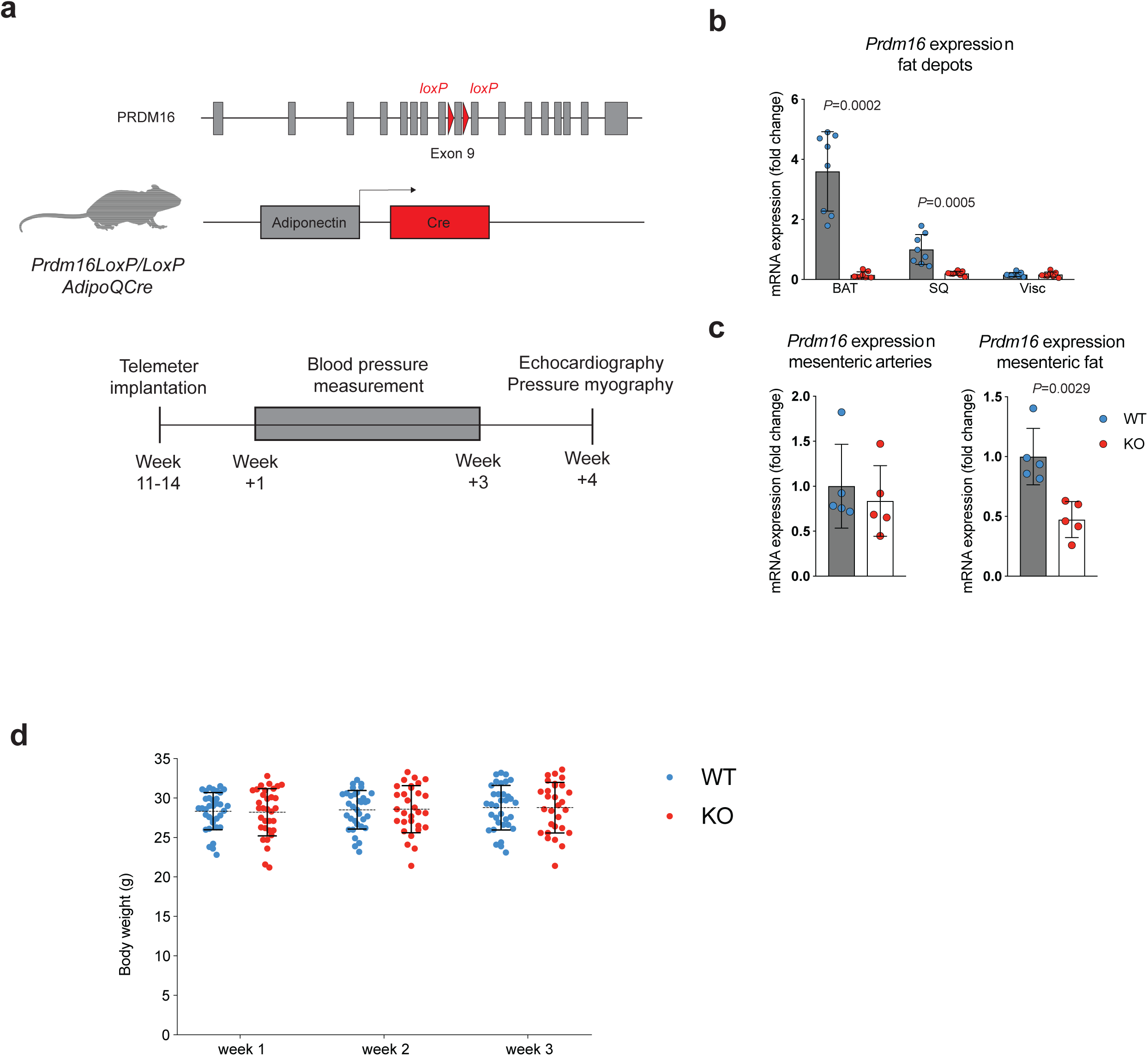
**a**, The loxP sites flanking exon 9 and the adiponectin driven Cre-recombinase that were used to create the *Prdm16* knockout (KO) mouse model are shown. Radio telemetry devices to measure blood pressure were implanted in mice between the age of 11 and 14 weeks and measurements were started 1 week after implantation for 3 weeks. Pressure myography and echocardiography were performed at the end of the study. **b**, Expression of *Prdm16* in brown adipose tissue (BAT), subcutaneous adipose tissue (SQ) and visceral adipose tissue (Visc) was measured by quantitative PCR (qPCR) and compared between KO (n=8) and littermate control wildtype animals (WT; n=8). Groups were compared using Mann-Whitney U test for non-normally distributed variables (BAT) and Student’s t-test for normally distributed variables (SQ and Visc). Bar depict means, error bars are s.e.m. All *P*-values are two-sided. **c**, Expression of *Prdm16* in mesenteric arteries and adjacent mesenteric adipose tissue was measured by quantitative PCR (qPCR) and compared between KO (n=4 for mesenteric arteries’ n=5 for mesenteric fat) and littermate control wildtype animals (WT; n=4 for mesenteric arteries’ n=5 for mesenteric fat). Groups were compared using Student’s t-test. Bar depict means, error bars are s.e.m. All *P*-values are two-sided. **d**, Comparison of body weights between WT and KO animals (both n=35 at the beginning of the study). Groups were compared using a mixed effects model with post-hoc analysis for individual time points.

**Extended Data Fig. 5.**
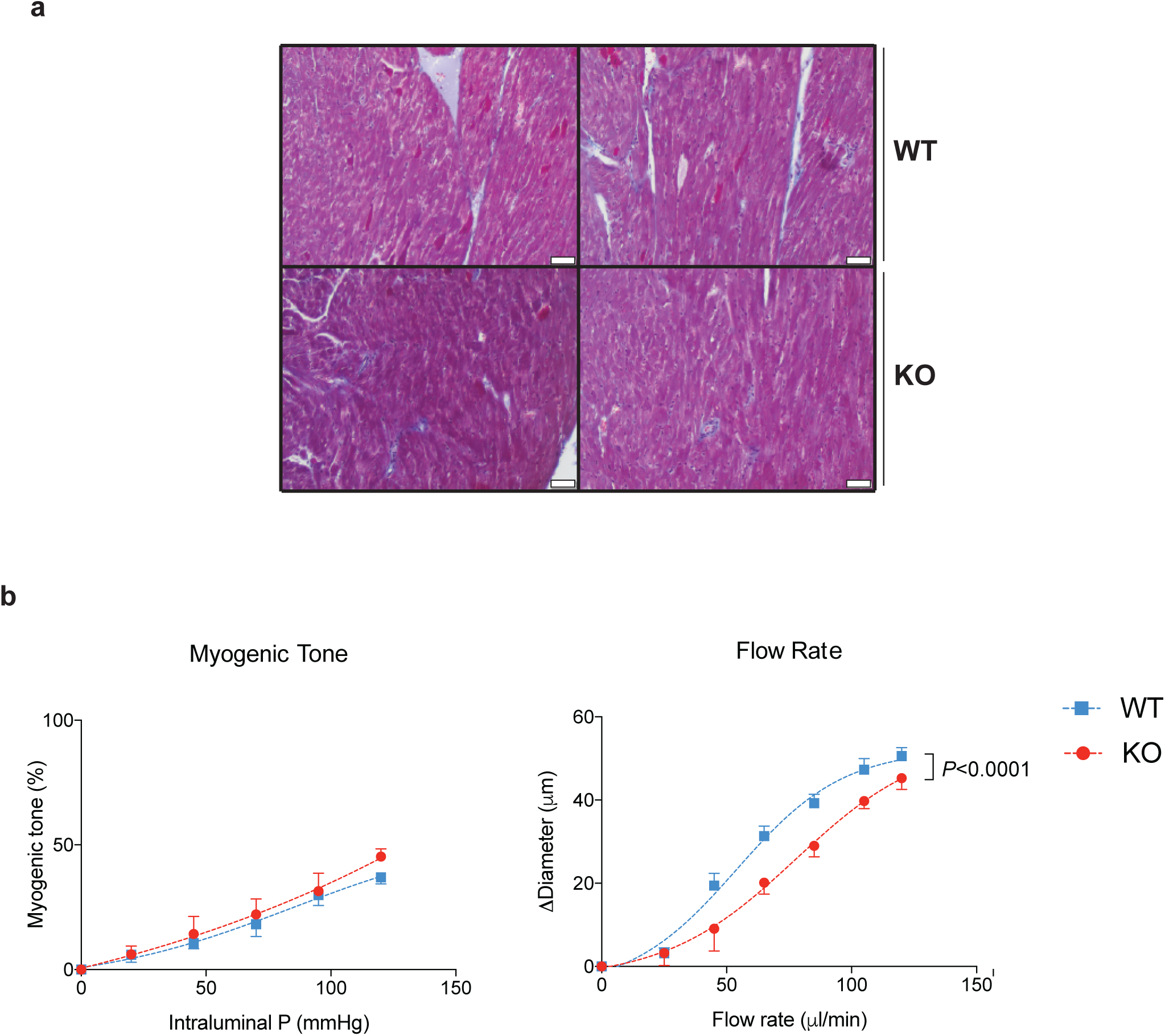
**a**, Sections of the left ventricle were stained with Masson’s trichrome stain to compare fat specific *Prdm16* knockout (KO) and littermate control (wildtype; WT) animals. Representative image, n=5 per group. Scale bar is 50 µm. **b**, Mesenteric arteries from KO and WT were harvested, cleaned of perivascular fat and used for pressure myography (n=3 per group). Groups were compared using two-way analysis of variance (ANOVA). Dots represent means, error bars s.e.m. All *P*-values are two-sided.

## References

1. Rosen, E.D. & Spiegelman, B.M. What we talk about when we talk about fat. Cell 156, 20–44 (2014).

2. Bartelt, A., et al. Brown adipose tissue activity controls triglyceride clearance. Nat. Med. 17, 200–205 (2011).

3. Lowell, B.B., et al. Development of obesity in transgenic mice after genetic ablation of brown adipose tissue. Nature 366, 740–742 (1993).

4. Cohen, P. & Spiegelman, B.M. Brown and Beige Fat: Molecular Parts of a Thermogenic Machine. Diabetes 64, 2346–2351 (2015).

5. Yeung, H.W., et al. Patterns of (18)F-FDG uptake in adipose tissue and muscle: a potential source of false-positives for PET. J. Nucl. Med. 44, 1789–1796 (2003).

6. Cohade, C., Osman, M., Pannu, H.K. & Wahl, R.L. Uptake in supraclavicular area fat (“USA-Fat”): description on 18F-FDG PET/CT. J. Nucl. Med. 44, 170–176 (2003).

7. van Marken Lichtenbelt, W.D., et al. Cold-activated brown adipose tissue in healthy men. N. Engl. J. Med. 360, 1500–1508 (2009).

8. Virtanen, K.A., et al. Functional brown adipose tissue in healthy adults. N. Engl. J. Med. 360, 1518–1525 (2009).

9. Cypess, A.M., et al. Identification and importance of brown adipose tissue in adult humans. N. Engl. J. Med. 360, 1509–1517 (2009).

10. Cronin, C.G., et al. Brown fat at PET/CT: correlation with patient characteristics. Radiology 263, 836–842 (2012).

11. Steinberg, J.D., Vogel, W. & Vegt, E. Factors influencing brown fat activation in FDG PET/CT: a retrospective analysis of 15,000+ cases. Br. J. Radiol. 90, 20170093 (2017).

12. Brendle, C., et al. Correlation of Brown Adipose Tissue with Other Body Fat Compartments and Patient Characteristics: A Retrospective Analysis in a Large Patient Cohort Using PET/CT. Acad. Radiol. 25, 102–110 (2018).

13. Ouellet, V., et al. Brown adipose tissue oxidative metabolism contributes to energy expenditure during acute cold exposure in humans. J. Clin. Invest. 122, 545–552 (2012).

14. Yoneshiro, T., et al. Brown adipose tissue, whole-body energy expenditure, and thermogenesis in healthy adult men. Obesity (Silver Spring) 19, 13–16 (2011).

15. Orava, J., et al. Different metabolic responses of human brown adipose tissue to activation by cold and insulin. Cell Metab. 14, 272–279 (2011).

16. Cohade, C., Mourtzikos, K.A. & Wahl, R.L. “USA-Fat”: prevalence is related to ambient outdoor temperature-evaluation with 18F-FDG PET/CT. J. Nucl. Med. 44, 1267–1270 (2003).

17. Kir, S. & Spiegelman, B.M. Cachexia & Brown Fat: A Burning Issue in Cancer. Trends in cancer 2, 461–463 (2016).

18. Cao, Q., et al. A pilot study of FDG PET/CT detects a link between brown adipose tissue and breast cancer. BMC Cancer 14, 126 (2014).

19. Fujii, T., et al. Implication of atypical supraclavicular F18-fluorodeoxyglucose uptake in patients with breast cancer: Association between brown adipose tissue and breast cancer. Oncology letters 14, 7025–7030 (2017).

20. Villarroya, F., Cereijo, R., Villarroya, J. & Giralt, M. Brown adipose tissue as a secretory organ. Nat Rev Endocrinol 13, 26–35 (2017).

21. Choi, C.H.J. & Cohen, P. Adipose crosstalk with other cell types in health and disease. Exp. Cell Res. 360, 6–11 (2017).

22. Wang, G.X., Zhao, X.Y. & Lin, J.D. The brown fat secretome: metabolic functions beyond thermogenesis. Trends Endocrinol Metab 26, 231–237 (2015).

23. Shinoda, K., et al. Genetic and functional characterization of clonally derived adult human brown adipocytes. Nat. Med. 21, 389–394 (2015).

24. Cohen, P., et al. Ablation of PRDM16 and beige adipose causes metabolic dysfunction and a subcutaneous to visceral fat switch. Cell 156, 304–316 (2014).

25. Liu, C., et al. Meta-analysis identifies common and rare variants influencing blood pressure and overlapping with metabolic trait loci. Nat. Genet. 48, 1162–1170 (2016).

26. Ward, Z.J., et al. Projected U.S. State-Level Prevalence of Adult Obesity and Severe Obesity. N. Engl. J. Med. 381, 2440–2450 (2019).

27. Cypess, A.M. & Kahn, C.R. Brown fat as a therapy for obesity and diabetes. Curr Opin Endocrinol Diabetes Obes 17, 143–149 (2010).

28. Blondin, D.P., et al. Dietary fatty acid metabolism of brown adipose tissue in cold-acclimated men. Nat Commun 8, 14146 (2017).

29. Chondronikola, M., et al. Brown Adipose Tissue Activation Is Linked to Distinct Systemic Effects on Lipid Metabolism in Humans. Cell Metab. 23, 1200–1206 (2016).

30. Yoneshiro, T., et al. Recruited brown adipose tissue as an antiobesity agent in humans. J. Clin. Invest. 123, 3404–3408 (2013).

31. Hanssen, M.J., et al. Short-term cold acclimation improves insulin sensitivity in patients with type 2 diabetes mellitus. Nat. Med. 21, 863–865 (2015).

32. Liu, X., et al. Brown adipose tissue transplantation improves whole-body energy metabolism. Cell Res. 23, 851–854 (2013).

33. Feldmann, H.M., Golozoubova, V., Cannon, B. & Nedergaard, J. UCP1 ablation induces obesity and abolishes diet-induced thermogenesis in mice exempt from thermal stress by living at thermoneutrality. Cell Metab. 9, 203–209 (2009).

34. Alcala, M., et al. Mechanisms of Impaired Brown Adipose Tissue Recruitment in Obesity. Front Physiol 10, 94 (2019).

35. Langenberg, C. & Lotta, L.A. Genomic insights into the causes of type 2 diabetes. Lancet 391, 2463–2474 (2018).

36. Yarnell, J.W., et al. Fibrinogen, viscosity, and white blood cell count are major risk factors for ischemic heart disease. The Caerphilly and Speedwell collaborative heart disease studies. Circulation 83, 836–844 (1991).

37. Friedman, G.D., Klatsky, A.L. & Siegelaub, A.B. The leukocyte count as a predictor of myocardial infarction. N. Engl. J. Med. 290, 1275–1278 (1974).

38. Ensrud, K. & Grimm, R.H., Jr. The white blood cell count and risk for coronary heart disease. Am. Heart J. 124, 207–213 (1992).

39. von Elm, E., et al. The Strengthening the Reporting of Observational Studies in Epidemiology (STROBE) statement: guidelines for reporting observational studies. Lancet 370, 1453–1457 (2007).

40. McGavigan, A.K., et al. Vertical sleeve gastrectomy reduces blood pressure and hypothalamic endoplasmic reticulum stress in mice. Disease models & mechanisms 10, 235–243 (2017).

41. Cantalupo, A., et al. Nogo-B regulates endothelial sphingolipid homeostasis to control vascular function and blood pressure. Nat. Med. 21, 1028–1037 (2015).

